# *Hydra*, a model system for deciphering the mechanisms of aging and resistance to aging

**DOI:** 10.1101/155804

**Authors:** Quentin Schenkelaars, Szymon Tomczyk, Yvan Wenger, Kazadi Ekundayo, Victor Girard, Wanda Buzgariu, Steve Austad, Brigitte Galliot

## Abstract

The freshwater cnidarian polyp named *Hydra*, which can be mass-cultured in the laboratory, is characterized by a highly dynamic homeostasis with a continuous self-renewal of its three adult stem cell populations, the epithelial stem cells from the epidermis, the epithelial stem cells from the gastrodermis, and the multipotent interstitial stem cells, which provide cells of the nervous system, gland cells and germ cells. Two unusual features characterize these stem cells that cannot replace each other, they all avoid G1 to pause in G2, and the two epithelial populations are concomitantly multifunctional and stem cells. *H. vulgaris* that does not show any signs of aging over the years, resists to weeks of starvation and adapts to the loss of neurogenesis, providing a unique model system to study the resistance to aging. By contrast some strains of a distinct species named *H. oligactis* undergo a rapid aging process when undergoing gametogenesis or when placed in stress conditions. The aging phenotype is characterized by the rapid loss of somatic interstitial stem cells, the progressive reduction in epithelial stem cell self-renewal, the loss of regeneration, the disorganization of the neuro-muscular system, the loss of the feeding behavior, and the death of all animals within about three months. We review here the possible mechanisms that help *H. vulgaris* to sustain stem cell self-renewal and thus bypass aging processes. For this, FoxO seems to act as a pleiotropic actor, regulating stem cell proliferation, stress response and apoptosis. In *H. oligactis,* the regulation of the autophagy flux differs between aging-sensitive and aging-resistant animals, pointing to a key role for proteostasis in the maintenance of a large pool of active and plastic epithelial stem cells.

---

The dramatic increase in average life expectancy is part of a major transition in human societies of the 21th century and global health issues linked to aging already have major economic and social impacts. Novel model systems are needed to complement existing ones (1). As an obvious strength of the *Hydra* model when compared to the classical genetic models of aging, namely the fruit fly *Drosophila melanogaster* and the worm *Caenorhabditis elegans*, adult polyps maintain at all times a highly dynamic homeostasis, thanks to abundant stocks of adult stem cells that continuously cycle and thus contribute to the lack of senescence and to the maintenance of amazing regenerative capacities. *Hydra* polyps are freshwater animals that belong to Cnidaria, a sister phylum to Bilateria (**Fig. 1A**). As all eumetazoans, *Hydra* differentiate a gut and a nervous system. These animals, which exhibit a tube shape morphology with a ring of tentacles, a mouth opening at the oral pole and a basal disc at the aboral one, are organized as two myoepithelial layers, epidermis outside and gastrodermis inside, separated by a thick acellular collagenous layer named mesoglea (**Fig. 1B)**. A nervous system made of sensory-motor neurons, ganglia neurons and mechanosensory cells named nematocytes (or cnidocytes) spreads across their body, densely packed at the apical and basal extremities, providing robust neuro-muscular activities and sophisticated behaviors (see in (2). The central part of the animal consists in epithelial and interstitial stem cells (ESCs and ISCs respectively) that continuously divide, while the extremities are predominantly made of cells that no longer divide.

**Figure 1:**
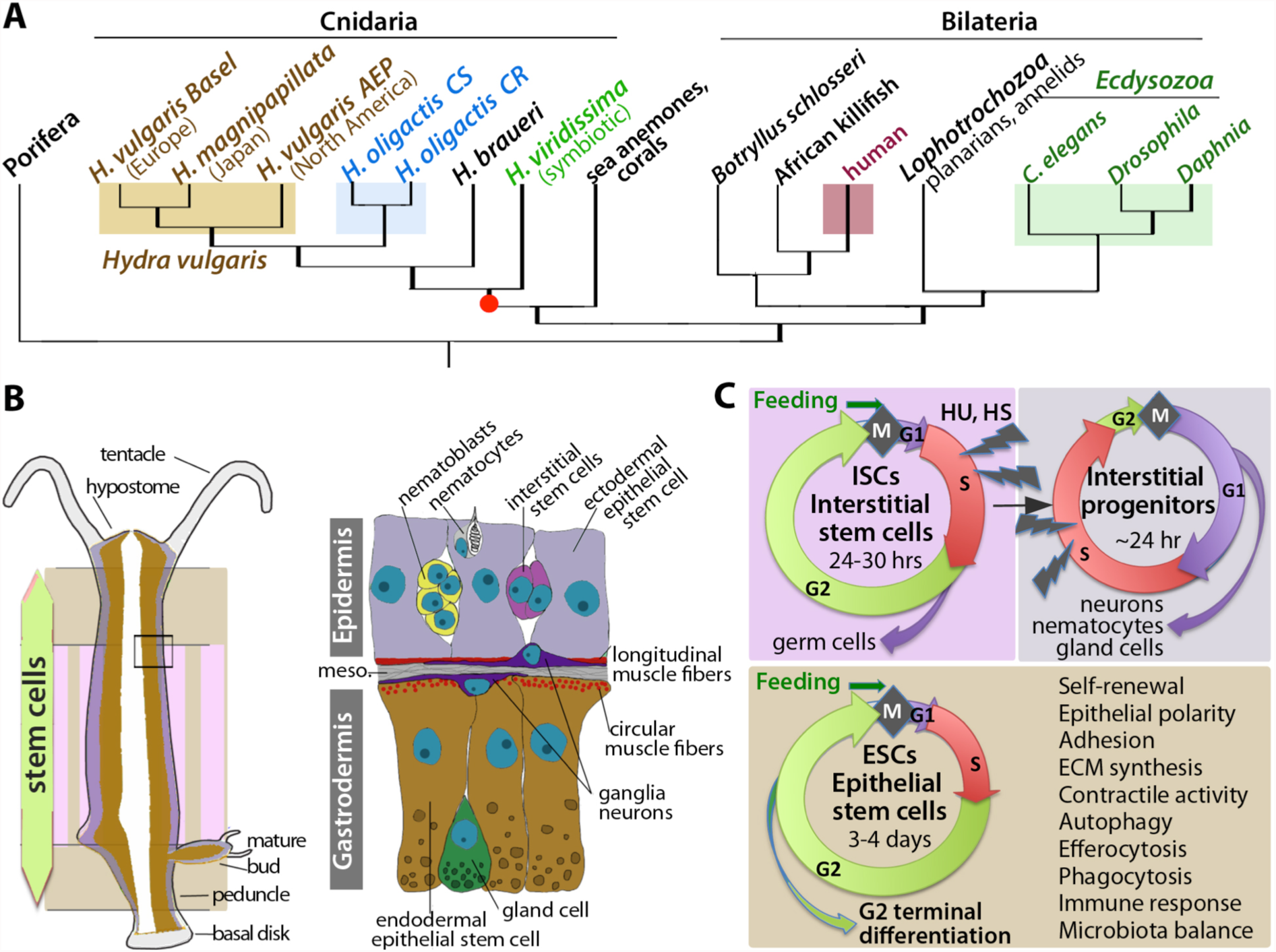
The *Hydra* model system. **(A)** Phylogenetic position of *Hydra* among metazoans. Note the sister group position of cnidarians to bilaterians and the close relationship between the various strains of *H. vulgaris*, and the two *H. oligactis* strains Cold Resistant (CR), Cold Sensitive (CS) used in aging studies. **(B)** Anatomy of *Hydra* polyp and tissue organization. Two distinct epithelial populations build the epidermal (purple) and gastrodermal (brown) layers. ESCs distribute along the body column (beige background) while ISCs, spread across the epidermis, are restricted to the central part of the animal (pink-beige background). The regions made of cells that no longer divide are colored light grey (tentacles, basal disk). The square indicates the enlarged region on the right. **(C)** Cell cycling behaviors of ESCs, ISCs and interstitial progenitors. ESCs are multifunctional cells, whose functions are listed on the right. ISCs and interstitial progenitors that cycle faster, are dramatically affected by colchicine (col), hydroxyurea (HU) and heat shock (HS) treatments (HS treatments are only efficient in the thermosensitive strain *Hv_sf1*).

## *Hydra vulgaris,* a slow- or no-aging animal

Evidences for low senescence in *Hydra* have accumulated since the pioneering study of Paul Brien in the mid 20th century (3). Indeed Brien nicely demonstrated that polyps of the *H. vulgaris (Hv,* named “*attenuata*” at that time) and *H. viridis* species maintained at 18°C exhibit stable budding and gametogenesis properties over several years (**Fig. 2)**. Fourty years later, Daniel Martinez monitored the mortality rates of four independent cohorts of animals over four years and confirmed these findings. He showed that despite a decrease in budding that he associates to environmental conditions, *Hv* polyps remained fit over this period, able to undergo gametogenesis, without exhibiting any obvious signs of senescence and basically no mortality (4). Martinez also pointed to the unusual longevity of *Hydra* polyps considering the age of first reproduction, two parameters that are correlated in most species. Although the origin of budding decline is debatable in the Martinez study (5), independent studies confirmed that *Hydra* are long-lived organisms with very low senescence over the years. Indeed, *Hv* polyps can be kept up to eight years (or more than 41 years for clonal culture) with a constant and very low annual probability of death (<1.5%) (6, 7). The view that emerged from these studies is that asexual as well as sexual *Hv* polyps do not age and thus represent a unique model system to investigate how an animal resists to aging.

**Figure 2:**
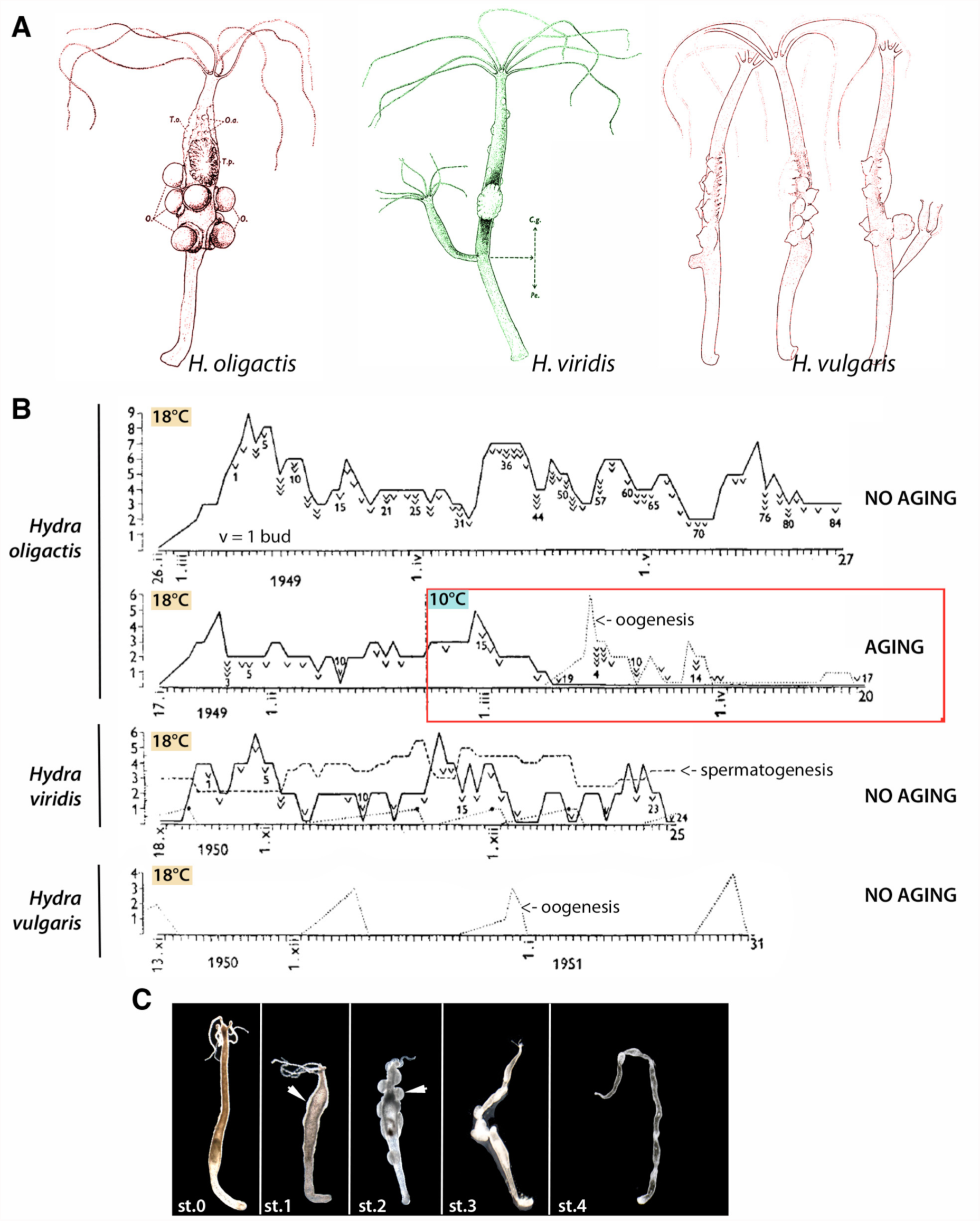
Discovery of *Hydra* aging by Paul Brien (1953) **(A)** Anatomies of a female *H. oligactis* (left), a hermaphrodite *H. viridis* (middle) and a male *H. vulgaris* (right) undergoing gametogenesis (after Figures 11-13, Brien, 1953). Cg: gastric column, O: oocytes, O.O: aborted eggs, Pe: peduncle, T.o: oocyte patch. **(B)** Individual records of budding and gametogenesis monitored in individual *H. oligactis, H. viridis* and *H. vulgaris* polyps (after Figure 21, Brien 1953). In each graph, budding is represented by a continuous line (each “v” indicates bud detachment), oogenesis by a dotted line (each detached oocyte represented as a small “v”), and spermatogenesis by a dashed line. Number of buds, oocytes, testes are indicated on the y axis, and time (days) on the x axis. Note that budding and spermatogenesis are continuous while oogenesis occurs as successive bursts. For *H. oligactis*, one polyp maintained at 18°C over three months shows a continuous budding, which persisted for 4 years at the same pace without any gametogenetic event (not shown). A second animal, transferred from 18°C to 10°C 35 days after birth, rapidly stopped budding after transfer while oogenesis was induced. Two months after transfer, Brien reports that the animal is “exhausted after having produced 17 eggs”. For *H. viridis* one polyp monitored over 10 weeks, produced 24 buds, 4 oocytes and multiple testes (the number of testes should be multiplied by 2). For *H. vulgaris*, one polyp was monitored over 11 weeks, undergoing 4 oogenetic events, each of them producing several oocytes. **(C)** Staging of the aging process in *Ho_CS* polyps from asexual healthy animal (stage 0), initiation of testis formation (arrow, stage 1), testes maturation and apical degeneration (arrow, stage 2), testis degeneration, loss of axial polarity and stenosis of the body column (stage 3), tissue loss and animal degeneration (stage 4).

## *Hydra oligactis,* a model for inducible aging

The situation is different when considering polyps from the closely related species *Hydra oligactis* (*Ho*). Indeed Paul Brien who wanted to investigate how animals from various *Hydra* species deal with a concomitant sexual and asexual reproduction, found that *Ho* animals transferred to 10°C, exhibit a rapid and dramatic reduction of their budding rate while undergoing sexual differentiation. He noticed that animals become “exhausted” after producing a series of oocytes (**Fig. 2B**) (3). This finding was confirmed by Lynne Littlefield in 1991 (8) and 15 years later more systematically investigated by Yoshida et al who described in details the aging phenotype (**Fig. 2C**), the morphological changes, the progressive alterations of the physiological functions such as food capturing and contractility, the similar mortality curves recorded in male and female colonies, but also the disorganization of the actin fibers and the dramatic reduction in ISCs (9). Interestingly, this *Ho* strain that we now name “cold-sensitive” (*Ho_CS*) is not representative of all *Ho* polyps as a closely related strain exhibits a “cold-resistant” (*Ho_CR*) behavior, i.e. undergoing sexual differentiation and transiently losing its somatic ISCs in the weeks following cold transfer, but surviving this “crisis” period and remaining fit over the following year when maintained at 10°C or transferred back to 18°C (10). Thus the two distinct *Ho_CS* and *Ho_CR* strains provide potent experimental models to get insights into the mechanisms that support aging and resistance to aging.

## Evolutionary conservation of the human aging gene families in *Hydra*

If we consider the 305 human aging genes listed on GenAge, one finds 73.1% orthologs or closely-related genes in the nematode *C. elegans*, 77.7% in the fruit fly *D. melanogaster*, 83% in *H. vulgaris* and 67.5% shared between the four species (**Fig. 3**). Indeed the analysis of the *H. vulgaris* genome (11) completed with transcriptomic analyses performed on intact *Hv_Basel* strain (12, 13) show that numerous human aging genes were lost in model organisms commonly used for aging studies (nematode, fruit fly), but conserved and expressed in *Hydra*, strengthening the value of this model to get insights into processes leading to aging and resistance to aging (**Fig. 3**). A series of complementary approaches were designed to quantify through RNA-seq transcriptomics the expression profile of each gene along the body column at five distinct positions (*spatial RNA-seq* (14)), in the three stem cell populations FACS-sorted from transgenic lines expressing GFP in one or the other stem cell population (*cell-type RNAseq* (15-17)), after elimination of ISCs by drugs or heat-shock (*epithelial plasticity RNA-seq* (17)), during *regeneration* (14, 18) and during *aging* in the *Ho_CS* and *Ho_CR* strains maintained for several weeks at 18°C or 10°C (Tomczyk et al. in preparation) (**Fig. 4A** and not shown, soon available online via HydrATLAS). In addition proteomic analyses were performed during regeneration (18) and aging (Tomczyk et al. in preparation). Such precise and systematic reports of genetic regulations and proteomic variations are a first step towards understanding the molecular mechanisms that underlie low senescence in *H. vulgaris* or aging in *Ho_CS*. According to (19) nine hallmarks define and contribute to aging in mammals: 1) stem cell exhaustion, 2) cellular senescence, 3) altered intercellular communication 3) deregulated nutrient sensing, 5) mitochondrial dysfunction, 6) genomic instability, 7) telomere attrition, 8) epigenetic alterations, 9) loss of proteostasis. We propose to discuss here how *Hydra* polyps may bypass some of these processes to defy death.

**Figure 3:**
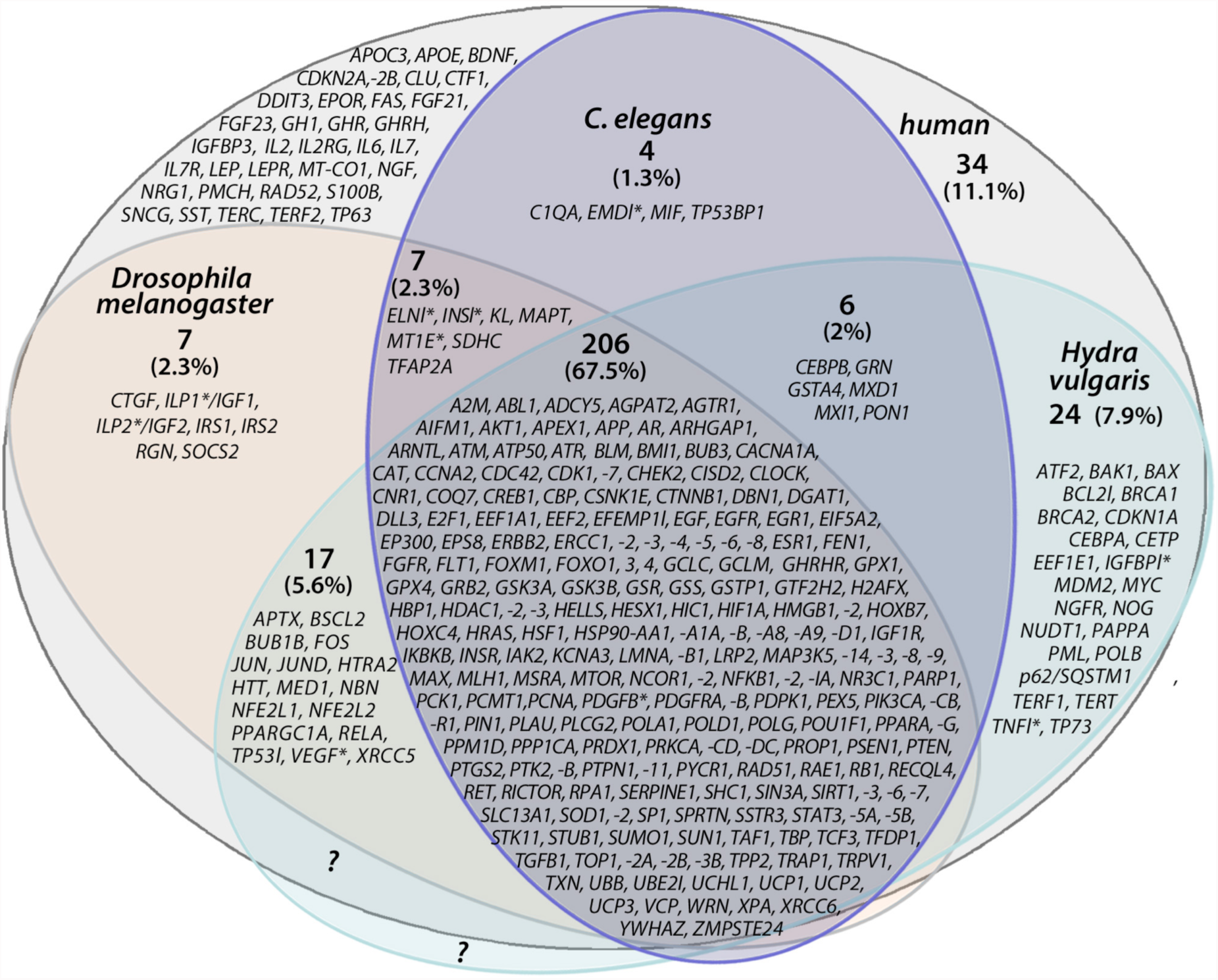
Venn diagram showing the distribution of 305 human aging gene families in *Drosophila melanogaster*, *Caenorhabditis elegans* and *Hydra vulgaris*. The deduced protein sequences of 305 human aging genes (grey background) listed in the database GenAge (genomics.senescence.info/genes/) were blasted on the NCBI and Swissprot/TrEMBL (uniprot.org) databases to identify the *Drosophila* (beige background), *C. elegans* (purple background) and *Hydra* (blue background) related sequences. Besthit sequences showing an e-value lower than 1^e^-10 were retrieved except few cases indicated with an asterisk where the e-value is >1^e^-10. Note that 24 gene families present in *H. vulgaris* were lost in *Drosophila* or *C. elegans*, while 34 human proteins do not have any counterparts in *Drosophila, C. elegans* or *Hydra*. Question marks indicate novel aging genes that might be discovered in *Hydra*, either conserved across evolution and present in human, or as taxon-specific innovations.

## Bypassing stem cell exhaustion

Stem cells continuously cycle in the *Hydra* body column while differentiated cells are predominantly found at the extremities, where epithelial cells are replaced within a week. Three distinct populations of stem cells populate *Hydra*: the unipotent epithelial stem cells (ESCs) of epidermis, the ESCs of the gastrodermis, and the multipotent interstitial stem cells (ISCs) that are spread across the epidermal layer (20-25). These three populations are distinct and cannot replace each other. However all stem cells in *Hydra* exhibit a rather unconventional cycling behavior: they pause in G2 and avoid G1 (**Fig. 1C**) (26). ESCs and ISCs show some striking differences as epithelial cells largely differentiate in G2 without traversing mitosis, forming thus terminally differentiated 4n cells (27), while ISCs produce progenitors that differentiate when reaching the G1 phase. ESCs fulfill the criteria of stem cells, i.e. self-renewing every three or four days when located in the body column, and terminally differentiating as soon as they cross sharp boundaries at each extremity of the body column, apical towards the head region, and basal at the aboral side (26, 28). More recently a pool of slow cycling stem cells, representing about 2% of each population, was identified, arrested in G2 for at least four weeks in case of ESCs (29).

All ESCs exhibit a low sensitivity to environmental conditions (starvation, stress) and are quite unconventional stem cells: they are unipotent and remain multifunctional at all times, continuously expressing for instance myoepithelial properties (**Fig. 1C)** (16). By comparison, the multipotent ISCs exhibit more classical stem cell features: located in the epidermal layer of the central region of the body column, they display a typical stem cell morphology, cycling every 24-30 hours, rapidly traversing G1 when cycling and pausing in G2, undergoing terminal differentiation as gland cells, nematocytes, nerve cells after a final mitosis in the body column or when reaching the extremities (**Fig. 1C**) (20, 24, 30). This spatially-restricted pattern of stem cell cycling suggests that *Hydra* polyps can be considered as indefinitely young in their central body column, and undergo aging at the extremities where cells get sloughed off and regularly replaced. All available information indicates that stem cell exhaustion does not exist in *H. vulgaris* strains maintained in favorable conditions (moderate temperature, regular feeding, absence of toxic substances).

How stem cells indefinitely keep cycling in the body column without genetic, epigenetic or metabolic alterations is a mystery. The self-renewal activity of stem cells is regulated by their own density and by the signals emitted between the various stem cells or by the differentiated cells (23). Two Myc transcription factors, one restricted to the somatic interstitial lineage (*Myc1*) and the second expressed in all stem cell populations (*Myc2*) are candidate regulators of the choice between self-renewal and differentiation (31-33). Indeed silencing *Myc1* expression or inhibiting Myc1 activity promotes proliferation of ISCs and their subsequent differentiation, suggesting that Myc1 is required to maintain the stemness of ISCs (32). Two independent transcriptomic analyses of the FACS-sorted stem cell populations identified the FoxO transcription factor as highly expressed by all stem cells in *Hydra* (15, 17) (see **Fig. 4B**). Functional assays indicate that FoxO acts as a key regulator of stem cell maintenance (34): when over-expressed in the interstitial lineage FoxO enhances stem cell proliferation and induces the expression of stem cell markers in differentiated cells, while its knockdown enhances cell differentiation, reduces animal growth and modifies innate immunity, but does not seem to induce an aging. It was proposed that the ancestral function of FoxO would be devoted to stem cell renewal, which in *Hydra* supports asexual reproduction, regeneration and immortality (35).

**Figure 4:**
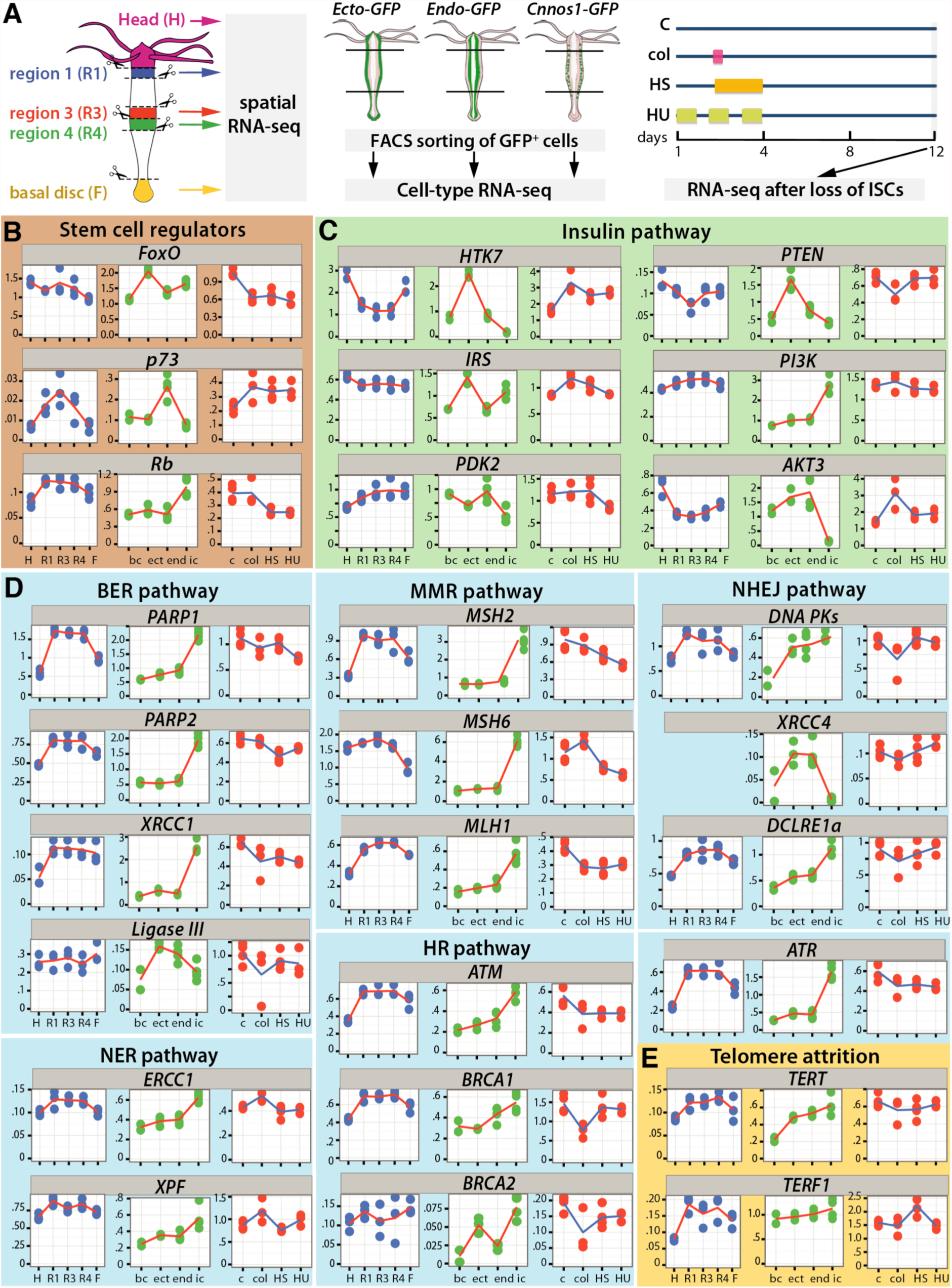
RNA-seq expression profiles of *Hydra* orthologs to human aging genes. **(A)** Procedures to identify the repertoire of genes expressed at different positions along the body axis in *Hv_Jussy* strain (spatial RNA-seq), in the different stem cell populations of *Hv_AEP* transgenic strains after FACS sorting (cell-type RNA-seq), after the loss of ISCs and neurogenesis upon transient colchicine (col), hydroxyurea (HU) or heatshock in *Hv_Sf1* strain. **(B-E)** Spatial (blue dots), cell-type (green dots) and ISC loss (red dots) RNA-seq profiles of orthologs to human aging genes involved in stem cell regulation (B), insulin pathway (C), DNA repair (D), telomere attrition (E). Each condition was tested in three biological replicates; bc: body column, ect: ectodermal ESCs, end: endodermal ESCs, ic: ISCs.

## No evidences for cellular senescence in *Hydra*

So far the presence and the role of cellular senescence is unknown in *Hydra*. Senescent cells are characterized by cell cycle arrest and thus, epithelial senescent cells should be detected outside the body column. β-galactosidase activity that is higher in senescent cells can be detected in *Hydra*, predominantly in the gastrodermis that contains predominantly cycling cells, measured at higher basal levels in *Ho_CS* than in *Hv_Basel*, enhanced upon feeding but also upon oxidative stress (Girard, unpublished). These preliminary results suggest a link between β-galactosidase activity and the efficiency of stress response, which is lower in *Ho_CS* (see below). In terms of known molecular markers of senescence, the cyclin-dependent kinase inhibitor CDKN2A (p16) that arrests cell cycling, does not have known counterpart in *Hydra*, *Drosophila*, or *C. elegans*. However the retinoblastoma gene RB1 that is well conserved across phyla, is expressed in all stem cells in *Hydra*, similarly to the transcription factors E2F1 and FoxO, whose interaction prevents activation of FoxO target genes and induces senescence in mammalian cells (36) (**Fig. 3, Fig. 4B**). Their respective role in regulating cell cycle arrest in *Hydra* needs to be tested.

## Bypassing deregulated nutrient sensing

### The Insulin, FoxO signaling pathway in *Hydra*

Genetic screens performed in *C. elegans* 30 years ago provided the first evidences of a genetic control of aging. The transcription factor Daf-16 (*foxO* ortholog) was identified as a central regulator of longevity, driving lifespan extension when the activity of the insulin/insulin-like growth factor (IGF) signaling pathway is low (37, 38). Indeed, mutations leading to a reduced activity of the insulin receptor Daf-2 increase worm lifespan, while the knockdown of *daf-16* reduces their lifespan. Briefly, when the activity of the insulin pathway is low, Daf-2 induces the phosphorylation of Daf-16 through AGE-1 and AKT, leading Daf-16 degradation (FoxO off). In contrast, when the insulin pathway is not activated, Daf-16 is no longer phosphorylated, can translocate to the nucleus where it activates a plethora of genes (FoxO on) that promote longevity by inducing resistance to oxidative stress, apoptosis, cell cycle arrest, DNA repair, differentiation. The pro-aging function of the insulin/IGF pathway and the pro-longevity function of Daf16/FoxO are shared between worms, *Drosophila* and mammals (39, 40).

In *Hydra* first evidences that the insulin/IGF pathway is functional came with the identification of HTK7, a receptor similar to the vertebrate insulin/IGF receptors, and the fact that polyps exposed to bovine insulin upregulate cell proliferation (41). Three insulin-like peptides (ILPs) were identified in the “Hydra peptide project” and in the genome (42). As *HTK7*, all three *ILP* genes are predominantly expressed by the ectodermal ESCs (**Fig 4C** and not shown). The genes encoding the kinases PKB/AKT (43), PDK2, Pi3K and its antagonizer PTEN are also heavily expressed in ESCs, an expression that persists when ISCs are eliminated (**Fig 4C**). As anticipated, the phosphorylated status of FoxO regulates its cytoplasmic versus nuclear localization at least in interstitital cells where Pi3K inhibition in a transgenic line overexpressing FoxO-GFP pushes FoxO-GFP to the nucleus (44). In a separate study, Lasi et al. showed that overexpressing FoxO-GFP in epithelial cells leads to apoptosis, an effect that is rescued by co-expressing *Hydra* ILP-1. This result implies that FoxO activity is under the control of the insulin/IGF pathway in epithelial cells (45). Bridge and coll. produced a transgenic line that overexpresses FoxO-GFP in interstitial cells and did not detect any pro-apoptotic effect of FoxO-GFP and no change in FoxO-GFP cellular localization upon starvation (44). These results might suggest that FoxO activity is differently regulated in the epithelial and interstitial lineages, and confirm a nutrient sensitivity restricted to ESCs in *Hydra*. An additional role of FoxO in the response to stress reported by (44) is discussed below.

### The autophagy and TOR pathways in *Hydra*

*Hydra* morphology and budding capacity is highly dependent on the nutrient conditions; animals rapidly stop budding upon food shortage and progressively reduce their size, but remain fit and survive weeks of starvation, exhibiting a very low mortality at least until 70 days of starvation (WB, unpublished). By contrast, heavily fed animals show an “obesity” phenotype, losing their typical apical-basal pattern and becoming highly heteromorphic (46, 47). As in most organisms, non-selective autophagy also named macro-autophagy is activated upon starvation in *Hydra,* a process that efficiently recycles cellular components in order to cope the absence of nutrients (48, 49). Double-membrane vacuoles named autophagosomes form in ESCs of starved animals, incorporating the lipidated form of an ubiquitin-like protein named LC3, and sequestering endosomes and mitochondria that get degraded once autophagosomes fuse with lysosomes. This dynamic autophagy flux can be pharmacologically modulated, by exposing starved polyps to Wortmannin, an inhibitor of autophagy entry, or to Bafilomycin that prevents lysosomal fusion. These findings highlight once more that in *Hydra,* ESCs are nutrient-sensitive. However the level of autophagy needs to be tightly balanced, as excessive autophagy dramatically impacts *Hydra* health as observed in animals knocked-down for the serine protease inhibitor Kazal1 (50). In such animals gigantic autophagosomes progressively form in ESCs, rapidly altering the ability of animals to survive body amputation, and progressively leading to animal death in the absence of injury. This phenotype mimics a pancreatitis autophagy phenotype observed in newborn mice knocked-out for *SPINK3*, or in patients harboring a mutated *SPINK1* gene, both genes being *Kazal*related. These studies indicate that autophagy plays an essential role in the environmental adaptation of *Hydra* (47, 50).

The evolutionary conserved nutrient-sensing pathway TOR (Target of Rapamycin) (51) is also well conserved in *Hydra* (e.g. *RAGs, TORC1, TORC2, Raptor, LST8, Ulk1/2*), predominantly expressed by the ESCs (16, 49). Rapamycin treatment, which mimics low nutrient conditions by inhibiting mTOR, acts as an autophagy inducer and indeed induces autophagosome formation in ectodermal ESCs of fed animals (48, 49). All together these data indicate that both the autophagy and the TOR pathways are efficient nutrient sensors in *Hydra.* Both pathways might actually play a key role in the aging phenotype identified in *Ho_CS,* as *Ho_CS* and *Ho_CR* polyps exhibit important differences in terms of autophagy when starved at 18°C or after transfer to 10°C. In addition Rapamycin treatment delays aging in *Ho_CS* (Tomczyk et al. in preparation).

## Bypassing mitochondrial dysfunction

The role of mitochondrial reactive oxygen species (ROS) during aging was recently reconsidered, rather assessed as a stress response to aging processes than a causal process that initiates aging (52). Not much is known on the role of mitochondria in the maintenance of *Hydra* homeostasis. Upon injury, mitochondrial superoxide is immediately produced by the endodermal ESCs, and this immediate wave of ROS signaling is likely beneficial for *Hydra*, triggering apoptosis of the most death-sensitive cells in the vicinity, but also cytoprotection via activation of genes encoding proteasome components, innate immune system regulators and stress proteins (14). FoxO plays an important role in the resistance to stress by upregulating the expression of genes encoding enzymes that detoxify damaging radicals (39). In *Hydra*, FoxO likely contributes to the stress response as evidenced by the JNK-dependent nuclear translocation of FoxO-GFP observed in heat-shocked animals (44, 53). Interestingly this regulation of FoxO appears similar in the epithelial and interstitial cell types of *H. vulgaris*. However the potential to adapt to heat shock (thermotolerance) is clearly different between *H. vulgaris* and *H.oligactis*, at the phenotypic level but also at the molecular level as *Hv* but not *Ho* rapidly produces the HSP70 protein (54), a difference explained by a very low stability of *hsp70* mRNAs in *Ho* (55). These results might reveal a more general deficiency in the ability to respond to stress in aging *Ho_CS*. Further studies will tell us whether accumulation of oxidative stress contributes to aging in *Ho_CS* and how FoxO contributes to face stress in *Hv, Ho_CR* and *Ho_CS* polyps.

## Bypassing genomic instability

Because DNA is the focal point where genetic information is stored, its integrity and stability are required for the maintenance of cell life and the transmission of error-free genome to the next generation (56). However, DNA is sensitive to stress-induced metabolic changes such as the production of highly reactive oxygen radicals and to environmental factors including radiation, which both induce DNA damages. To face these issues, pathways aimed at preventing damages or at promoting DNA repair are constantly activated (57). Briefly, preventing DNA damages mainly consists in the direct removal of spontaneous addition of methyl groups by MGMT, AlkB or MPG, while DNA repair relies on five major pathways, base excision repair (BER), nucleotide excision repair (NER) and mismatch repair (MMR), which act on single strand damages, bulky lesions and mismatches respectively, while non-homologous end joining (NHEJ) and homologous recombination (HR) pathways repair double strand breaks.

Little is known about the function and the regulation of these pathways in *Hydra*, even though most components were characterized in *Hydra* (**Fig. 3**). The *Hydra* Xeroderma Pigmentosum group A ortholog (XPA) is highly similar to human XPA, encoding the ERCC1 helix-hairpin-helix motif and a nuclear localization signal (58). Similarly the function of Xeroderma Pigmentosum group F (XPF), which encodes an ERCC4 endonuclease domain and is predominantly expressed in ISCs, is likely conserved in *Hydra* (59). Concerning the BER pathway, genes encoding the Flap endonuclease 1 (Fen1), XRCC1, Polymerase-β (Polβ) and Ligase3 were identified in genomic and transcriptomic analyses in *Hydra* (11, 13) as well as in the jellyfish *Aurelia aurita* (60). This pathway shows a highly variable evolution, with Polβ expressed in several arthropods (*Tribolium castaneum, Halyomorpha halys*, *Putella xylostella*) but absent in the fruit fly or in nematodes (**Fig. 3**). Such independent secondary losses imply that Polβ may play a redundant function in the BER pathway.

To get insights into the DNA repair processes that might be active in *Hydra*, we show here a compilation of the RNAseq profiles of some key genes involved in DNA damage sensing (*ATR*), BER (i.e. *PARP1, PARP2, XRCC1, Ligase 3*), NER (i.e. *ERCC1, XPF*), MMR (i.e. *MSH2, MSH6, MLH1*), HR (i.e. *ATM, BRCA1, BRCA2*) and NHEJ (*DNA PKs, DCLR1a, XRCC4*) pathways (**Fig. 4D**). *DNA-PKs* is expressed in all stem cell lineages, *Ligase 3* and *XRCC4* are predominantly expressed in ESCs, while all the other genes are expressed at higher levels in ISCs, consistent with the fact that these fast-cycling cells that spend less time in G2-phase than ESCs, likely require highest levels of DNA repair. Moreover, ISCs also provide germ cells, which require an efficient DNA repair machinery to maintain their genome integrity and preserve the next generation from incorrect genetic information. Although quantitative transcriptomics bring some hints on the potential role of DNA repair in *Hydra* stem cells, their function as a support to regeneration and low senescence remains to be elucidated. Also, since DNA repair mechanisms contribute to cell cycle arrest and apoptosis (61), it would be of interest to investigate their putative role in the dramatic changes that affect stem cells in aging *Ho_CS.*

## Bypassing telomere attrition

The progressive shortening of telomeres in somatic cells contributes to the aging process since replicative senescence or apoptosis occur when the telomere length falls below a certain threshold (Hayflick limit) (62). As a consequence telomere length measured in early life can be used as a reliable predictive value of lifespan in various organisms (63). In dividing cells, telomere shortening is actively counterbalanced by the telomerase reverse transcriptase (TERT) that protects chromosomes from telomere attrition by adding small DNA repeats at chromosome extremities (64). Interestingly in choanoflagellates, porifers, cnidarians including *Hydra*, and humans, TERT adds the same DNA motif TTAGGG, suggesting some conservation of an ancestral motif across evolution, while this motif diverged in nematodes (TTAGGC) and arthropods (TTAGG). Accordingly, TERT activity can be used to estimate the level of cellular immortality. While TERT activity becomes undetectable in differentiated cells, it is maintained high in germ cells, and reactivated in cancer stem cells, explaining why these cells continuously divide without undergoing senescence (65). High levels of telomerase activity were also measured in somatic cells of two marine demosponges (66) as well as in somatic cells of asexual but not sexual planarians (67). Interestingly, these species do not seem to age and possess robust regenerative capacities, supporting the hypothesis that low senescence is linked to high TERT activity.

In *Hydra* little is known about telomere attrition and TERT activity. Although Traut and collaborators failed to detect TERT activity in *Hydra,* they showed that TERT from another cnidarian, the jellyfish *Aurelia aurita,* is capable to add repeats to an artificial substrate (64). The transcriptomic data produced in our laboratory on each stem cell population of the body column and in different locations along the body axis (17) show that the telomeric repeat-binding factor 1 (*Terf1*) that negatively regulates telomere length is similarly expressed in ASCs and in differentiated cells of the body column (**Fig. 4E**). However *Tert* expression is lower at the extremities of the animal, and consistently with this finding, higher in the three stem cell populations than in the differentiated cells of the body column. These results support the idea that TERT activity is enhanced in *Hydra* somatic stem cells, protecting the genome from telomere attrition. Thus, promoting elongation of telomeres may take part in the processes that allow *Hydra* stem cells to conspicuously divide without any sign of exhaustion, but functional studies are needed to evidence the role of TERT in maintaining stem cell integrity and low senescence in *Hydra*.

## Bypassing epigenetic alterations

The activity of transcription factors or transcriptional complexes that regulate gene expression through gene regulatory elements (GRE), is modulated by epigenetic changes that consist in DNA methylation and histone modifications (68). Thanks to genome wide ChIP-seq analyses, the Technau lab identified a similar distribution and abundance of GREs in the sea anemone *Nematostella vectensis* and in bilaterians (69). For instance, they identified high levels of H3K4me3 in the vicinity of transcriptional start sites (TSS) and the mutual exclusion between CpG methylation and H3K4me3, consistent with their opposed roles on gene expression. Furthermore, the predicted binding sites of the transcriptional co-factor p300 correspond to enhancer elements enriched in similar regions than in *Drosophila*. Thus, same principles apply for complex gene regulation and chromatin modifications in cnidarians and bilaterians, most likely already present in their last common ancestor. Similarly, the *Hydra* genome is methylated (2.7%) while those from *Drosophila* and *C. elegans* lack DNA methylation marks (70). Although this finding certainly reflects the small genome size in *D. melanogaster* and *C. elegans*, it remains true that these features of cnidarian genomes provide a framework to study epigenetic modifications and make *Hydra* a relevant model system to investigate the role of epigenetic alterations in the aging processes.

The Polycomb repressive complexes (PRC1, PRC2) are potent chromatin modulators in bilaterians (71). Suppression subtractive hybridization and *in situ* hybridization revealed that the *Hydra* ortholog of the PRC2 histone methlytransferase EED (*<u>e</u>mbryonic <u>e</u>ctoderm <u>d</u>evelopment*, *HyEED*) is express in all embryonic cells at early stages, becoming restricted to ISCs, nematoblast and spermatogonia in the body column in adult animals (72). The production of a transgenic line where HyEED is overexpressed as a chimeric HyEED-eGFP protein under the ubiquitous actin promoter, revealed that EED-eGFP is stable only in ISCs and differentiating nematoblasts and undetectable at the animal extremities, suggesting that EED-eGFP is degraded in differentiated cells (73). Also, EED-eGFP localization is both cytoplasmic and nuclear, suggesting that it binds chromatin. All these observations are consistent with a repressive role of PRC2 on the differentiation of stem cells or progenitors (74). Interestingly, when proteasome degradation is inhibited in these transgenic animals, HyEED-eGFP+ cells are detected in head and tentacles, supporting the hypothesis of proteasomal degradation in differentiated cells (73). Understanding how stem cell activity is controlled by PRCs thus appears as a promising trend for aging studies.

## Bypassing the loss of proteostasis

The research on diseases linked to aging has experienced an unprecedented advance over the past 20 years with the discovery that the onset of numerous diseases is linked to the loss of protein homeostasis (proteostasis), which is controlled by genetic pathways and biochemical processes largely conserved across evolution (75). Briefly, it was estimated that about 30% of the newly synthetized proteins are misfolded (76). To clear such proteins, healthy cells use two main pathways for their degradation, lysosomal proteases acting on autophagic vacuoles and the ubiquitin proteasomal system. When both pathways are overflowed, misfolded proteins readily accumulate as protein aggregates that contain proteins normally not bound to each other. Then secondary modifications such as ubiquitinylation and cross-linking take place providing highly reactive aggregates that recruit new proteins (75, 77). As an amplifying feedback loop, aggregates inhibit proteasomal clearance, leading to severe metabolic changes that damage cells (78). To degrade large aggregates, macroautophagy (refered here as autophagy) remains the only efficient process, a process named aggrephagy (79). The accumulation of aggregates and the inhibition of the proteasome seem to impact more dramatically cells that no longer divide, explaining the vulnerability of neurons as observed in neurodegenerative diseases, but also the vulnerability of epithelial cells in multiple organs, probing the development of cancers, age-linked myopathies, retinal dystrophies, cataract, arteriopathies.

The high cellular turnover that characterizes *Hydra* homeostasis certainly prevents the effect of aggregate formation by diluting aggregates throughout successive cell division. However, aggrephagy also exists in *Hydra* to get rid of aggregates and reduce the risks of impaired proteostasis (Tomczyk et al. in preparation). *Ho_CS* polyps are much more sensitive to proteasome inhibition than *Ho_CR* or *Hv* polyps (**Fig. 5A**), likely revealing the deficient compensatory autophagy in *Ho_CS* as shown by the accumulation of selective autophagy marker p62/SQSTM1. This deficient proteostasis dramatically impacts the self-renewal of ESC, contributing to the aging phenotype of *Ho_CS* animals. These promising results indicate that *Hydra* also provides a relevant model to dissect the mechanisms that regulate the cross-talk between proteostasis and stem cell activity.

**Figure 5:**
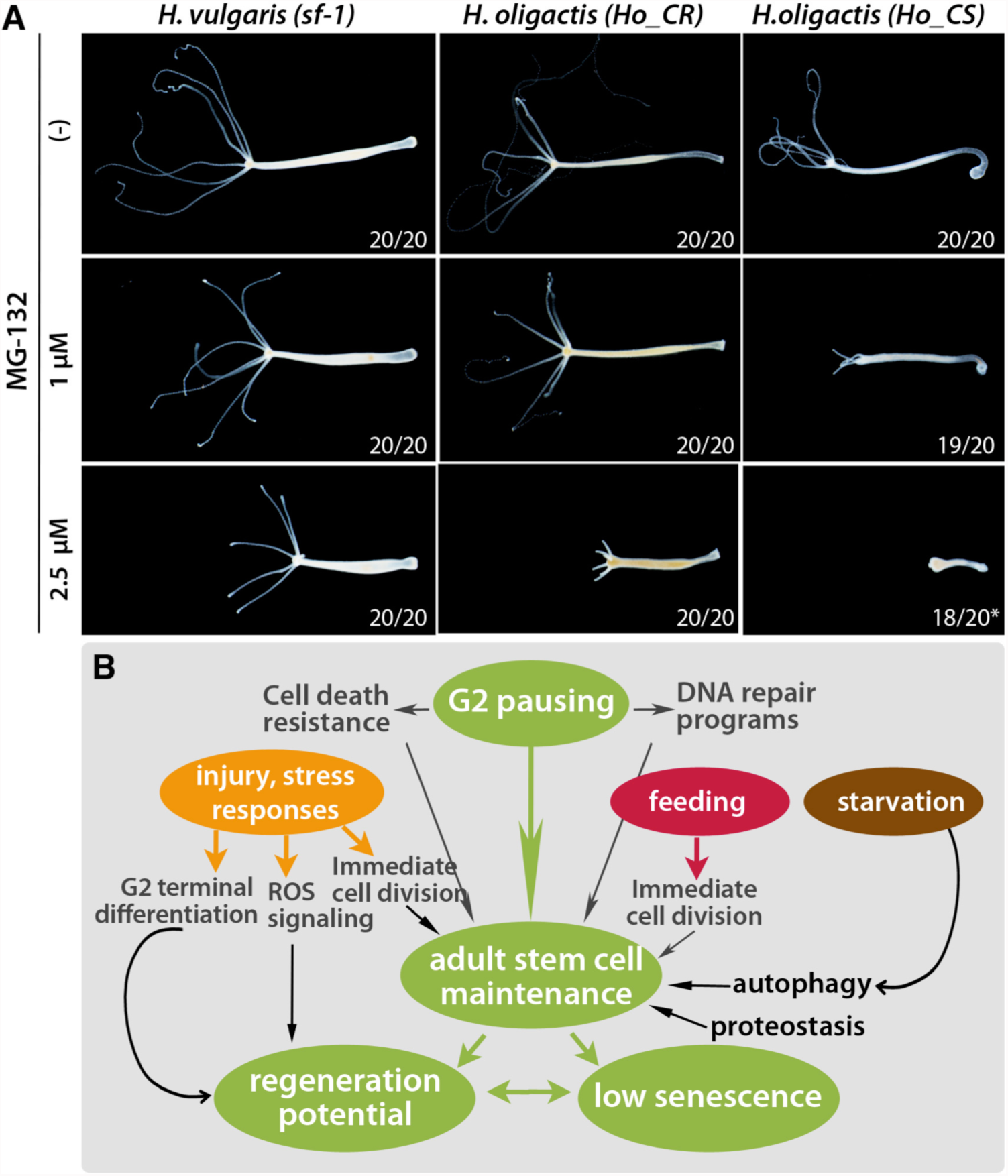
Sensitivity to proteostasis in *Hydra* and summary scheme. **(A)** Toxicity of a 3 day MG132 treatment on *Hv_SF1, Ho_CR* and *Ho_CS* animals maintained at 18°C starved during the experiment. Note the higher sensitivity of *Ho_CS* compared to *Ho_CR* or *Hv_sf1.* Asterisks indicate two dead animals. **(B)** Summary scheme of the cellular and molecular regulations that contribute or are supposed to contribute to the maintenance of active pools of stem cells in *Hydra*, a necessary condition for regeneration and slow aging.

## Conclusions & Perspectives

Genome-scale bioinformatic analyses have shown that most gene families and pathways involved in mammalian aging processes are expressed in *Hydra*. Biochemical assays, live imaging tools as well as genetic assays such as transgenesis or gene silencing just started to dissect how *H. vulgaris* resist to aging, and how environmental changes induce aging in *H. oligactis,* a process that leads to animal death within a short period (three months). Indeed some strains of *H. oligactis* are prone to undergo aging when the stock of somatic interstitial cells is depleted as a result of gametogenesis or environmental stress. In this context, the plasticity of epithelial stem cells appears deficient and self-renewal declines, seemingly as a result of combined deficient proteostasis, deficient autophagy and/or to oxidative stress. By contrast *H. vulgaris* seem to bypass critical processes that lead to animal death, maintaining unexhausted its pool of multifunctional ESCs that adapt to harsh environmental conditions such as prolonged starvation or loss of neurogenesis. A series of properties that maintain dynamic and unaltered the pool of stem cells, emerge as pro-longevity features in adult *Hydra* (**Fig. 5B**). These properties are linked to specificities of stem cell cycling such as G2 pausing that promotes DNA repair and cell death resistance, but also to the multifunctionality of ESCs, which readily adapt to environmental threats by mounting efficient autophagy and stress responses. Although phylogenetically distant from mammals, *Hydra* provides a novel model to trace the evolutionary-conserved basis of animal aging, to identify potential taxon-specific innovations that help animals resist to aging, and to screen for molecules that modify aging processes.

## Acknowledgements

This work was supported by the Swiss National Science Foundation (SNF 31003A_149630), the National Institute of Health (grant NIH-R01AG037962), the canton of Geneva (Switzerland). The authors thank T. Bosch (Kiel Germany) for kindly providing the AEP transgenic strains used to produce the cell-type RNA-seq data.

## REFERENCES

1. Austad SN. (2009). Is there a role for new invertebrate models for aging research? J Gerontol A Biol Sci Med Sci 64, 192-194.

2. Galliot B. (2012). Hydra, a fruitful model system for 270 years. Int J Dev Biol 56, 411-423.

3. Brien P. (1953). La Perennite Somatique. Biol Rev 28, 308-349.

4. Martinez DE. (1998). Mortality patterns suggest lack of senescence in hydra. Exp Gerontol 33, 217-225.

5. Estep PW. (2010). Declining asexual reproduction is suggestive of senescence in hydra: comment on Martinez, D., “Mortality patterns suggest lack of senescence in hydra.” Exp Gerontol 33, 217-25. Exp Gerontol 45, 645-646.

6. Jones OR, Scheuerlein A, Salguero-Gomez R, Camarda CG, Schaible R, Casper BB, Dahlgren JP, Ehrlen J, et al. (2014). Diversity of ageing across the tree of life. Nature 505, 169-173.

7. Schaible R, Scheuerlein A, Danko MJ, Gampe J, Martinez DE, Vaupel JW. (2015). Constant mortality and fertility over age in Hydra. Proc Natl Acad Sci U S A 112, 15701-15706.

8. Littlefield CL, Finkemeier C, Bode HR. (1991). Spermatogenesis in Hydra oligactis. II. How temperature controls the reciprocity of sexual and asexual reproduction. Dev Biol 146, 292-300.

9. Yoshida K, Fujisawa T, Hwang JS, Ikeo K, Gojobori T. (2006). Degeneration after sexual differentiation in hydra and its relevance to the evolution of aging. Gene 385, 64-70.

10. Tomczyk S, Fischer K, Austad S, Galliot B. (2015). Hydra, a powerful model for aging studies. Invertebr Reprod Dev 59, 11-16.

11. Chapman JA, Kirkness EF, Simakov O, Hampson SE, Mitros T, Weinmaier T, Rattei T, Balasubramanian PG, et al. (2010). The dynamic genome of Hydra. Nature 464, 592-596.

12. Wenger Y, Galliot B. (2013). Punctuated emergences of genetic and phenotypic innovations in eumetazoan, bilaterian, euteleostome, and hominidae ancestors. Genome Biol Evol 5, 1949-1968.

13. Wenger Y, Galliot B. (2013). RNAseq versus genome-predicted transcriptomes: a large population of novel transcripts identified in an Illumina-454 Hydra transcriptome. BMC genomics 14, 204.

14. Wenger Y, Buzgariu W, Reiter S, Galliot B. (2014). Injury-induced immune responses in Hydra. Semin Immunol 26, 277-294.

15. Hemmrich G, Khalturin K, Boehm AM, Puchert M, Anton-Erxleben F, Wittlieb J, Klostermeier UC, Rosenstiel P, et al. (2012). Molecular signatures of the three stem cell lineages in hydra and the emergence of stem cell function at the base of multicellularity. Mol Biol Evol 29, 3267-3280.

16. Buzgariu W, Al Haddad S, Tomczyk S, Wenger Y, Galliot B. (2015). Multi-functionality and plasticity characterize epithelial cells in Hydra. Tissue Barriers 3, e1068908.

17. Wenger Y, Buzgariu W, Galliot B. (2016). Loss of neurogenesis in Hydra leads to compensatory regulation of neurogenic and neurotransmission genes in epithelial cells. Philos Trans B 371, 20150040.

18. Petersen HO, Hoger SK, Looso M, Lengfeld T, Kuhn A, Warnken U, Nishimiya-Fujisawa C, Schnolzer M, et al. (2015). A Comprehensive Transcriptomic and Proteomic Analysis of Hydra Head Regeneration. Mol Biol Evol 32, 1928-1947.

19. Lopez-Otin C, Blasco MA, Partridge L, Serrano M, Kroemer G. (2013). The hallmarks of aging. Cell 153, 1194-1217.

20. Bode HR. (1996). The interstitial cell lineage of hydra: a stem cell system that arose early in evolution. J Cell Sci 109, 1155-1164.

21. Galliot B, Miljkovic-Licina M, de Rosa R, Chera S. (2006). Hydra, a niche for cell and developmental plasticity. Semin Cell Dev Biol 17, 492-502.

22. Watanabe H, Hoang VT, Mattner R, Holstein TW. (2009). Immortality and the base of multicellular life: Lessons from cnidarian stem cells. Semin Cell Dev Biol 20, 1114-1125.

23. Bosch TC, Anton-Erxleben F, Hemmrich G, Khalturin K. (2010). The Hydra polyp: nothing but an active stem cell community. Dev Growth Differ 52, 15-25.

24. David CN. (2012). Interstitial stem cells in Hydra: multipotency and decision-making. Int J Dev Biol 56, 489-497.

25. Hobmayer B, Jenewein M, Eder D, Eder MK, Glasauer S, Gufler S, Hartl M, Salvenmoser W. (2012). Stemness in Hydra - a current perspective. Int J Dev Biol 56, 509-517.

26. Buzgariu W, Crescenzi M, Galliot B. (2014). Robust G2 pausing of adult stem cells in Hydra. Differentiation 87, 83-99.

27. Dubel S, Schaller HC. (1990). Terminal differentiation of ectodermal epithelial stem cells of Hydra can occur in G2 without requiring mitosis or S phase. J Cell Biol 110, 939-945.

28. Bosch TC, David CN. (1984). Growth regulation in Hydra: relationship between epithelial cell cycle length and growth rate. Dev Biol 104, 161-171.

29. Govindasamy N, Murthy S, Ghanekar Y. (2014). Slow-cycling stem cells in hydra contribute to head regeneration. Biol Open 3, 1236-1244.

30. David CN, Plotnick I. (1980). Distribution of interstitial stem cells in Hydra. Dev Biol 76, 175-184.

31. Hartl M, Mitterstiller AM, Valovka T, Breuker K, Hobmayer B, Bister K. (2010). Stem cell-specific activation of an ancestral myc protooncogene with conserved basic functions in the early metazoan Hydra. Proc Natl Acad Sci U S A 107, 4051-4056.

32. Ambrosone A, Marchesano V, Tino A, Hobmayer B, Tortiglione C. (2012). Hymyc1 downregulation promotes stem cell proliferation in Hydra vulgaris. PLoS One 7, e30660.

33. Hartl M, Glasauer S, Valovka T, Breuker K, Hobmayer B, Bister K. (2014). Hydra myc2, a unique pre-bilaterian member of the myc gene family, is activated in cell proliferation and gametogenesis. Biol Open 3, 397-407.

34. Boehm AM, Khalturin K, Anton-Erxleben F, Hemmrich G, Klostermeier UC, Lopez-Quintero JA, Oberg HH, Puchert M, et al. (2012). FoxO is a critical regulator of stem cell maintenance in immortal Hydra. Proc Natl Acad Sci U S A 109, 19697-19702.

35. Schaible R, Sussman M. (2013). FOXO in aging: did evolutionary diversification of FOXO function distract it from prolonging life? Bioessays 35, 1101-1110.

36. Xie Q, Peng S, Tao L, Ruan H, Yang Y, Li TM, Adams U, Meng S, et al. (2014). E2F transcription factor 1 regulates cellular and organismal senescence by inhibiting Forkhead box O transcription factors. The Journal of biological chemistry 289, 34205-34213.

37. Friedman DB, Johnson TE. (1988). A mutation in the age-1 gene in Caenorhabditis elegans lengthens life and reduces hermaphrodite fertility. Genetics 118, 75-86.

38. Kenyon C, Chang J, Gensch E, Rudner A, Tabtiang R. (1993). A C. elegans mutant that lives twice as long as wild type. Nature 366, 461-464.

39. Greer EL, Brunet A. (2005). FOXO transcription factors at the interface between longevity and tumor suppression. Oncogene 24, 7410-7425.

40. Martins R, Lithgow GJ, Link W. (2016). Long live FOXO: unraveling the role of FOXO proteins in aging and longevity. Aging cell 15, 196-207.

41. Steele RE, Lieu P, Mai NH, Shenk MA, Sarras MP. (1996). Response to insulin and the expression pattern of a gene encoding an insulin receptor homologue suggest a role for an insulin-like molecule in regulating growth and patterning in *Hydra*. Dev Genes Evol 206, 247-259.

42. Fujisawa T, Hayakawa E. (2012). Peptide signaling in Hydra. Int J Dev Biol 56, 543-550.

43. Herold M, Cikala M, MacWilliams H, David CN, Bottger A. (2002). Cloning and characterisation of PKB and PRK homologs from Hydra and the evolution of the protein kinase family. Dev Genes Evol 212, 513-519.

44. Bridge D, Theofiles AG, Holler RL, Marcinkevicius E, Steele RE, Martinez DE. (2010). FoxO and stress responses in the cnidarian Hydra vulgaris. PLoS One 5, e11686.

45. Lasi M, David CN, Bottger A. (2010). Apoptosis in pre-Bilaterians: Hydra as a model. Apoptosis 15, 269-278.

46. Otto JJ, Campbell RD. (1977). Tissue economics of hydra: regulation of cell cycle, animal size and development by controlled feeding rates. J Cell Sci 28, 117-132.

47. Galliot B, Ghila L. (2010). Cell plasticity in homeostasis and regeneration. Mol Reprod Dev 77, 837-855.

48. Buzgariu W, Chera S, Galliot B. (2008). Methods to investigate autophagy during starvation and regeneration in hydra. Methods Enzymol 451, 409-437.

49. Chera S, Buzgariu W, Ghila L, Galliot B. (2009). Autophagy in Hydra: a response to starvation and stress in early animal evolution. Biochim Biophys Acta 1793, 1432-1443.

50. Chera S, de Rosa R, Miljkovic-Licina M, Dobretz K, Ghila L, Kaloulis K, Galliot B. (2006). Silencing of the hydra serine protease inhibitor Kazal1 gene mimics the human SPINK1 pancreatic phenotype. J Cell Sci 119, 846-857.

51. Loewith R, Hall MN. (2011). Target of rapamycin (TOR) in nutrient signaling and growth control. Genetics 189, 1177-1201.

52. Hekimi S, Lapointe J, Wen Y. (2011). Taking a “good” look at free radicals in the aging process. Trends Cell Biol 21, 569-576.

53. Martinez DE, Bridge D. (2012). Hydra, the everlasting embryo, confronts aging. Int J Dev Biol 56, 479-487.

54. Bosch TC, Krylow SM, Bode HR, Steele RE. (1988). Thermotolerance and synthesis of heat shock proteins: these responses are present in Hydra attenuata but absent in Hydra oligactis. Proc Natl Acad Sci U S A 85, 7927-7931.

55. Brennecke T, Gellner K, Bosch TC. (1998). The lack of a stress response in Hydra oligactis is due to reduced hsp70 mRNA stability. Eur J Biochem 255, 703-709.

56. Vijg J, Suh Y. (2013). Genome instability and aging. Annu Rev Physiol 75, 645-668.

57. Lombard DB, Chua KF, Mostoslavsky R, Franco S, Gostissa M, Alt FW. (2005). DNA repair, genome stability, and aging. Cell 120, 497-512.

58. Barve A, Ghaskadbi S, Ghaskadbi S. (2013). Structural and sequence similarities of hydra xeroderma pigmentosum A protein to human homolog suggest early evolution and conservation. Biomed Res Int 2013, 854745.

59. Barve A, Ghaskadbi S, Ghaskadbi S. (2013). Conservation of the nucleotide excision repair pathway: characterization of hydra Xeroderma Pigmentosum group F homolog. PLoS One 8, e61062.

60. Kodera H, Takeuchi R, Uchiyama Y, Takakusagi Y, Iwabata K, Miwa H, Hanzawa N, Sugawara F, et al. (2011). Characterization of marine X-family DNA polymerases and comparative analysis of base excision repair proteins. Biochem Biophys Res Commun 415, 193-199.

61. Branzei D, Foiani M. (2008). Regulation of DNA repair throughout the cell cycle. Nat Rev Mol Cell Biol 9, 297-308.

62. Hayflick L, Moorhead PS. (1961). The serial cultivation of human diploid cell strains. Exp Cell Res 25, 585-621.

63. Heidinger BJ, Blount JD, Boner W, Griffiths K, Metcalfe NB, Monaghan P. (2012). Telomere length in early life predicts lifespan. Proc Natl Acad Sci U S A 109, 1743-1748.

64. Traut W, Szczepanowski M, Vitkova M, Opitz C, Marec F, Zrzavy J. (2007). The telomere repeat motif of basal Metazoa. Chromosome Res 15, 371-382.

65. Shay JW, Wright WE. (2010). Telomeres and telomerase in normal and cancer stem cells. FEBS Lett 584, 3819-3825.

66. Koziol C, Borojevic R, Steffen R, Muller WE. (1998). Sponges (Porifera) model systems to study the shift from immortal to senescent somatic cells: the telomerase activity in somatic cells. Mech Ageing Dev 100, 107-120.

67. Tan TC, Rahman R, Jaber-Hijazi F, Felix DA, Chen C, Louis EJ, Aboobaker A. (2012). Telomere maintenance and telomerase activity are differentially regulated in asexual and sexual worms. Proc Natl Acad Sci U S A 109, 4209-4214.

68. Jaenisch R, Bird A. (2003). Epigenetic regulation of gene expression: how the genome integrates intrinsic and environmental signals. Nat Genet 33 Suppl, 245-254.

69. Schwaiger M, Schonauer A, Rendeiro AF, Pribitzer C, Schauer A, Gilles AF, Schinko JB, Renfer E, et al. (2014). Evolutionary conservation of the eumetazoan gene regulatory landscape. Genome Res 24, 639-650.

70. Hassel M, Cornelius MG, Vom Brocke J, Schmeiser HH. (2010). Total nucleotide analysis of Hydra DNA and RNA by MEKC with LIF detection and 32P-postlabeling. Electrophoresis 31, 299-302.

71. Blackledge NP, Rose NR, Klose RJ. (2015). Targeting Polycomb systems to regulate gene expression: modifications to a complex story. Nat Rev Mol Cell Biol 16, 643-649.

72. Genikhovich G, Kurn U, Hemmrich G, Bosch TC. (2006). Discovery of genes expressed in Hydra embryogenesis. Dev Biol 289, 466-481.

73. Khalturin K, Anton-Erxleben F, Milde S, Plotz C, Wittlieb J, Hemmrich G, Bosch TC. (2007). Transgenic stem cells in Hydra reveal an early evolutionary origin for key elements controlling self-renewal and differentiation. Dev Biol 309, 32-44.

74. Zoabi M, Sadeh R, de Bie P, Marquez VE, Ciechanover A. (2011). PRAJA1 is a ubiquitin ligase for the polycomb repressive complex 2 proteins. Biochem Biophys Res Commun 408, 393-398.

75. Balchin D, Hayer-Hartl M, Hartl FU. (2016). In vivo aspects of protein folding and quality control. Science 353, aac4354.

76. Schubert U, Anton LC, Gibbs J, Norbury CC, Yewdell JW, Bennink JR. (2000). Rapid degradation of a large fraction of newly synthesized proteins by proteasomes. Nature 404, 770-774.

77. Grune T, Jung T, Merker K, Davies KJ. (2004). Decreased proteolysis caused by protein aggregates, inclusion bodies, plaques, lipofuscin, ceroid, and ‘aggresomes’ during oxidative stress, aging, and disease. Int J Biochem Cell Biol 36, 2519-2530.

78. Bence NF, Sampat RM, Kopito RR. (2001). Impairment of the ubiquitin-proteasome system by protein aggregation. Science 292, 1552-1555.

79. Lamark T, Johansen T. (2012). Aggrephagy: selective disposal of protein aggregates by macroautophagy. Int J Cell Biol 2012, 736905.

